# Efficient and Rapid generation of anti-aging neural stem cells by direct conversion Fibroblasts with A Single microRNA

**DOI:** 10.1101/2021.08.09.452601

**Authors:** Yuanyuan Li, Jing Sun, Tingting Xu, Xiaobin Weng, Yanan Zhang, Theodore P. Rasmussen, Bo Dai, Yuesi Wang

## Abstract

Neural stem cells (NSCs) have great potential in the application of neurodegenerative disease therapy, drug screening and disease modeling. NSC can be generated by reprogramming from terminally differentiated cells with transcription factors or small molecules. However, current methods for producing NSCs involve the danger of integrating foreign genes into the genome and the problem of low efficiency. Here, we report an efficient method to generate NSCs from human skin-derived fibroblasts with microRNA (mir-302a) in 2-3 days. The induced NSCs (iNSCs) have more than 90% of purity. Their morphology is similar to regular NSCs, expressing key markers including Nestin, Pax6 and Sox2, and can be expanded for more than 20 passages in vitro. They can also differentiate into functional neuron progeny, astrocytes and oligodendrocytes as well. Those cells can elicit action potential, can be xeno-transplanted into the brain of immune-deficient mice, and can survive and differentiate in vivo without tumor formation. This study shows that a single part of pluripotency-inducing mir-302 cluster can drive fibroblasts reprogramming, providing a general platform for high-efficiency generation of individual-specific human NSCs for studies of neuron system development and regenerative cell therapy.

## Introduction

Neural stem cells (NSCs) have great potentials in cell therapy, drug screening, tissue engineering, and development research of neural degenerative diseases, stroke, and injury^[1]^. However, the limited cell resources and the limited number of NSCs hinder the research and clinical application of these cells. Cell reprogramming or conversion is promising to generate NSCs from terminally differentiated cells^[2]^. Induced Neural stem cells(iNSCs) can be obtained by transfecting somatic cells with pluripotent transcription factors such as Sox2 or Oct4, either alone or in combination with other transcription factors^[3–4]^. Neural stem cells can be generated by transfection with non-pluripotent transcription factors, such as BMi1, Zfp521, Ptf1, etc^[5–11]^. However, with the present techniques, the inefficiency, time-consuming, and low purity of iNSCs by direct conversion limit their potential applications in cell therapy.

To improve efficiency and reduce time consumption, the most frequently used approaches to generate iNSCs are through transcriptional factors (TFs) transfection with small molecules^[12–17]^. One study employed Sox2 to reprogram human skin fibroblasts into iNSCs in only four days. However, the study did not give a corresponding colony formation rate or reprogramming efficiency^[18]^. Despite these encouraging advances with TFs and small molecules, the existing protocols that use translational factor or small-molecule compounds to reprogram somatic cells into iNSCs are still tedious, requiring up to more than two weeks with relatively low reprogramming efficiency (0.08%-66%) and colonies formation efficiency ( 0.0002%-0.6% ) (Table S1). Time-consuming, low efficiencies, and paltry yields hinder the application of neural stem cells for cell therapy. More effective reprogramming strategies must be found to solve these problems.

It has been shown that the microRNAs-mediated direct conversion approach has a high efficiency ^[19–21]^. In previous studies, microRNA mir-302/367cluster has been shown to high efficiently reprogram human and mouse fibroblasts into iPSCs. The study specifically showed that mir-302a/b/c/d did not produce any iPSC colonies without mir-367 in murine fibroblasts^[22]^. However, the results of another paper showed that mir-302 clusters can reprogram human skin cancer cells into a pluripotent state of ES-like^[23]^. The increase in reprogramming efficiency produced with mir-302 clusters in those studies is still not sufficient. Moreover, the use of multiple components cannot account for the effect of individual components. Since multiple components together can have the ability to reprogram somatic cells, does a single component have a similar function to reprogram fibroblasts into NSCs? The study found that mir-302a was highly expressed in the NSC derived from H9 embryonic stem cells^[24]^. All these together lead us to hypothesize that mir-302a alone might be able to reprogram fibroblasts to have neural stem cell potential. To determine whether overexpression of a single component mir-302a of the mir-302 cluster could reprogram fibroblasts, we constructed lentiviral/adenovirus vectors, encoding the has-mir-302a(mir-302a) sequence , one component of the mir-302 cluster , and used it to reprogram human postnatal foreskin derived fibroblasts (HFFs, healthy donors, 26- and 29-year-old males) and Eyelid fibroblasts ( healthy donors , a 41-year-old female ).

To generate neural stem cells more rapidly and efficiently, we developed a reprogramming strategy combining microRNA, pharmaceutical molecules^[12, 25]^, and glass coverslips culture^[3]^ that is faster and more efficient than existing methods. Human fibroblasts were cultured on gelatin-coated glass coverslips in fibroblast culture medium in 24-well-plate overnight. On the second day, the medium was changed into a fresh medium containing has-mir-302a virus with polybrene, VPA, and Vitamin C 24 hours post-transfection, NSCs induction media were added into the medium. After overexpression of mir-302a in HFFs within 24 to 48 hours, their morphology changes quickly and form clusters (Figure 1a). Using a live cell workstation and inverted fluorescence microscope to observation , the first colony emerged as early as 13 hours (Figure 1b , Figure S1, Supporting Information). Three days after transfection with mir-302a ,the colony-forming rate of HFFs is as high as 1.1 % , then reached 2.29 % on day 5 and 8.2 % on day 7. Eyelid fibroblasts, similar to HFFs, can rapidly form neurospheres. The colony-forming rate of eyelid fibroblasts was as high as 3.04 % on day 3 (Figure 1c, Figure S,2,3,4, Supporting Information) ,showing the faster and more efficient than any currently available method for deriving neurospheres from human fibroblasts ( Table S1 ). Three days later, almost all the transfected cells formed neurospheres. The efficiency of reprogramming can be increased by adding different concentrations of VPA, VC, and polybrene. At the optimal combination of the three concentrations, GFP-tagged transfected cells showed the efficiency of cell reprogramming was up to 90 % ( Figure 1d ). It is more efficient and faster than existing methods of reprogramming into neural stem cells with transfection reprogramming ( Table S1 ).

**Figure 1.**
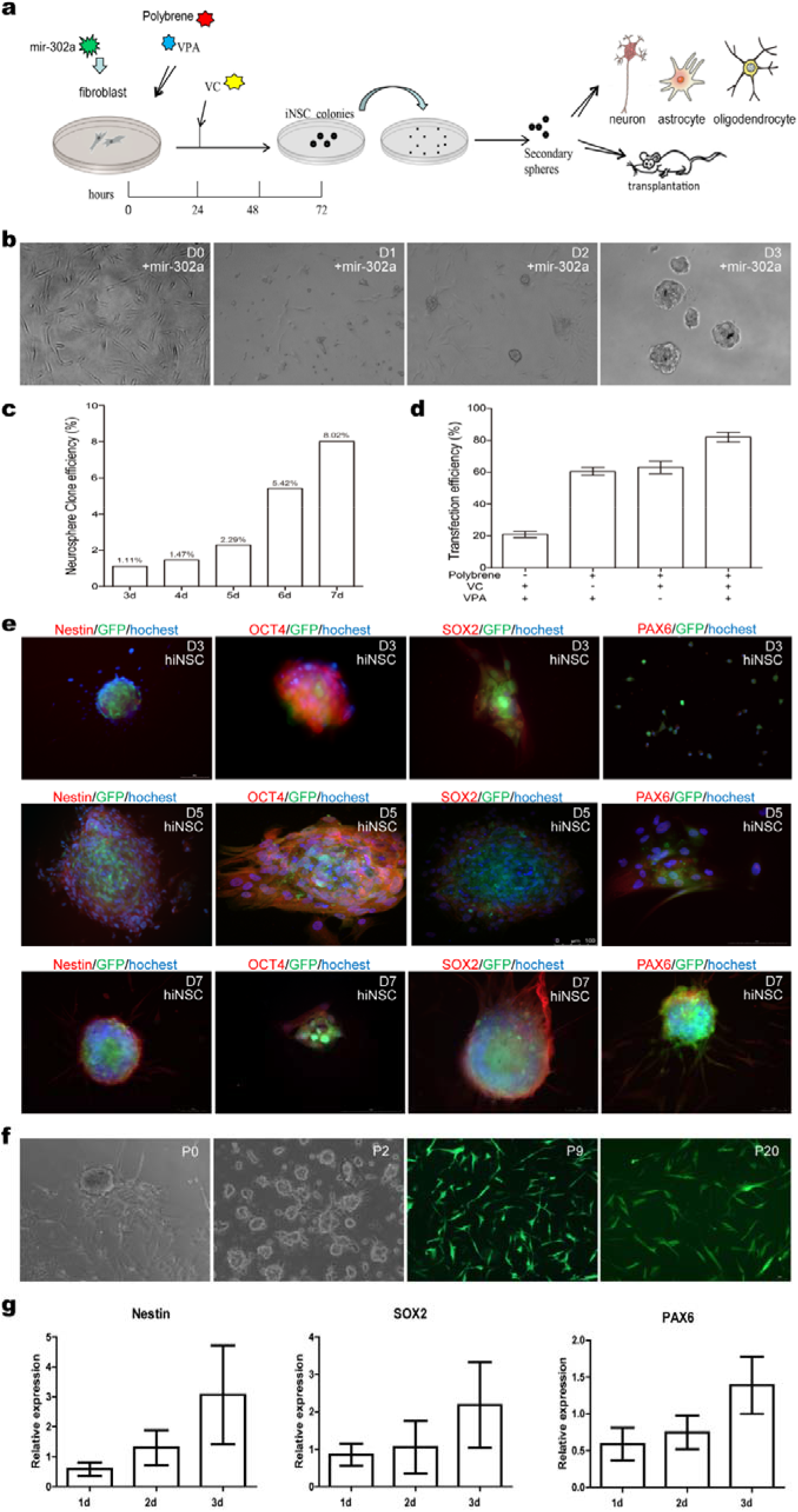
Generation and Characterization of iNSCs from Human Postnatal Foreskin-Derived Fibroblasts (HFFs). a) Schematic representation of the single miRNA mir-302a reprogramming process to generate neurospheres from human fibroblasts. b) Phase-contrast images of HFFs after overnight treatment with mir-302a virus vector in fibroblast medium.has-mir-302a-infected cells spontaneously formed neurosphere-like colonies on gelatin-coated glass coverslips from 1-3 days after infection. c)The colony formation efficiency of neurospheres generated by hiNSCs from human fibroblasts at 3,4,5,6, and 7 days after mir-302a transfection. d)Effect of the three small molecules used to increase transfection efficiency for reprogramming with has-mir-302a. e) Expression of Nestin, Oct4, Pax6 and Sox2 in hiNSC neurospheres at 3, 5, 7 days after mir-302a transfection. f) Neurospheres derived from mir-302a reprogrammed human fibroblasts can be expanded from P0 to P20 passages with adherent and suspension culture alternately. g) qRT-PCR reveals that hiNSCs express typical NSC markers. On day 1, typical NSC markers Nestin were expressed. On day 3, hiNSCs express typical NSC markers Nestin, Pax6, and Sox2.

To verify whether the obtained cells are neural stem cells, we use immunofluorescence to detect NSC cell surface markers Nestin, Sox2, and Pax6. Immunostaining results showed that the colonies’ expression of key NSC markers Nestin, Sox2, and Pax6 could be detected within 72 hours, with escalation gradually during prolonged culture( Figure 1e ). The results of immunofluorescence strikingly confirmed the close similarity between hiNSCs and the control human Ptf1a-induced iNSCs (Figure S5, Supporting Information). The appearance of cell surface markers indicates that the reprogrammed cells have the surface characteristics of neural stem cells.

To demonstrate the purity and yield of the iNSCs obtained, flow cytometry measurements of the purity and yield of iNSCs conversion showed that nearly 90% of surviving lentivirally infected fibroblasts are Nestin positive on day 3. Nestin-positive cells from these colonies were about 97% ( Figure S6, Supporting Information ), the results showing that the fibroblasts were converted into NSCs, revealing an unprecedented higher purity and yield of conversion.

To demonstrate that the obtained cells have the ability to self-renew, the induced neurospheres were collected, dissociated, and replated into NSC expansion media with suspension culture and monolayer culture. The new iNSCs secondary neurospheres rapidly formed starting from day 2. Suspension culture and monolayer experience are applied alternately to expand and purify neural stem cells. The iNSCs could be expanded and maintained stably for over ten months and more than 20 passages. The morphologies of iNSCs in passage 1, 2 , 9, and 20 were stable and homogenous, which were all Nestin , Sox2 and Pax6 positive (Figure 1g) , which indicating their self-renewal ability.

To prove the role of using glass slice culture to improve the efficiency of neural stem cell reprogramming compared to direct plate laying, we used the Real-time PCR technique to detect the gene expression of neural stem cells. Real-time quantitative PCR showed that Nestin, Pax6, and Sox2 were expressed on the first day of reprogramming HFFs into neural stem cells, and increased with time. Expression of Nestin, Pax6, and Sox2 of hiNSCs reprogrammed on glass slides was stronger than in cells directly attached to culture plates, and the formation of neurospheres was faster(Figure S7, Supporting Information). The protein expression of Oct4 in hiNSCs is weak and gradually disappeared within a short time, Oct4 expression in mir-302a-hiNSCs was transient(Figure S4,Supporting Information). This result is somewhat similar to that of Bag óJR et al results. h-iNSCs did not have a high expression of Oct4,Oct4 is not expressed after h-iNSC differentiation^[18]^.

To verify that mir-302a can reprogram not only human fibroblasts but also murine fibroblasts, mouse L929 cells are also rapidly and efficiently reprogrammed by mir-302a into neural stem cells (Figure S8, Supporting Information). On day 3, 80% of the cells rounded and clustered, with neurospheres forming. The expression of Nestin, Pax6, and Sox2 in the induced miNSCs was similar to that in the control mouse neural stem cell line SCR029. The method is also more efficient and faster than the existing methods of reprogramming mouse neural stem cells (Table S2). Overall, these data demonstrate that a single mir-302a is sufficient to rapidly and efficiently reprogram human and mouse fibroblasts into large numbers of neural stem cells.

We further evaluated whether the obtained neural stem cells can differentiate into neurons and glial cells. The developmental potential of iNSCs was assessed by their capacity to generate neurons, astrocytes, and oligodendrocytes. The method of spontaneous differentiation of neurons is to culture neural stem cells on a feeder layer for three days. After the withdrawal of the feeders, bFGF, and EGF, iNSCs differentiated into a large number of Tuj1 positive neurons within four days. After an additional 1 to 2 week culture, iNSCs differentiated into Tuj1 ,Map2 ,SYN ,NSE and NeuN positive neurons (Figure 2a-b). Quantitative examination at three weeks showed that 89% of the cells were Tuj1 positive (Figure 2e). Immunocytochemistry showed iNSC derived astrocytes were GFAP and Vimentin positive. Furthermore, hiNSCs can also differentiate into MBP and Olig2 positive cells during oligodendrocyte differentiation (Figure 2c-d). Under the astrocyte differentiation condition, 92% of the cells differentiated into GFAP positive astrocytes. In the oligodendrocyte differentiation medium, 94% of differentiated cells were MBP-positive (Figure 2e). The pivotal surface markers expression pattern is similar in iNSCs from different fibroblasts sources. These results strongly indicate that mir-302a-reprogrammed-iNSCs can give rise to multiple neuronal subtypes and show pluripotency.

**Figure 2.**
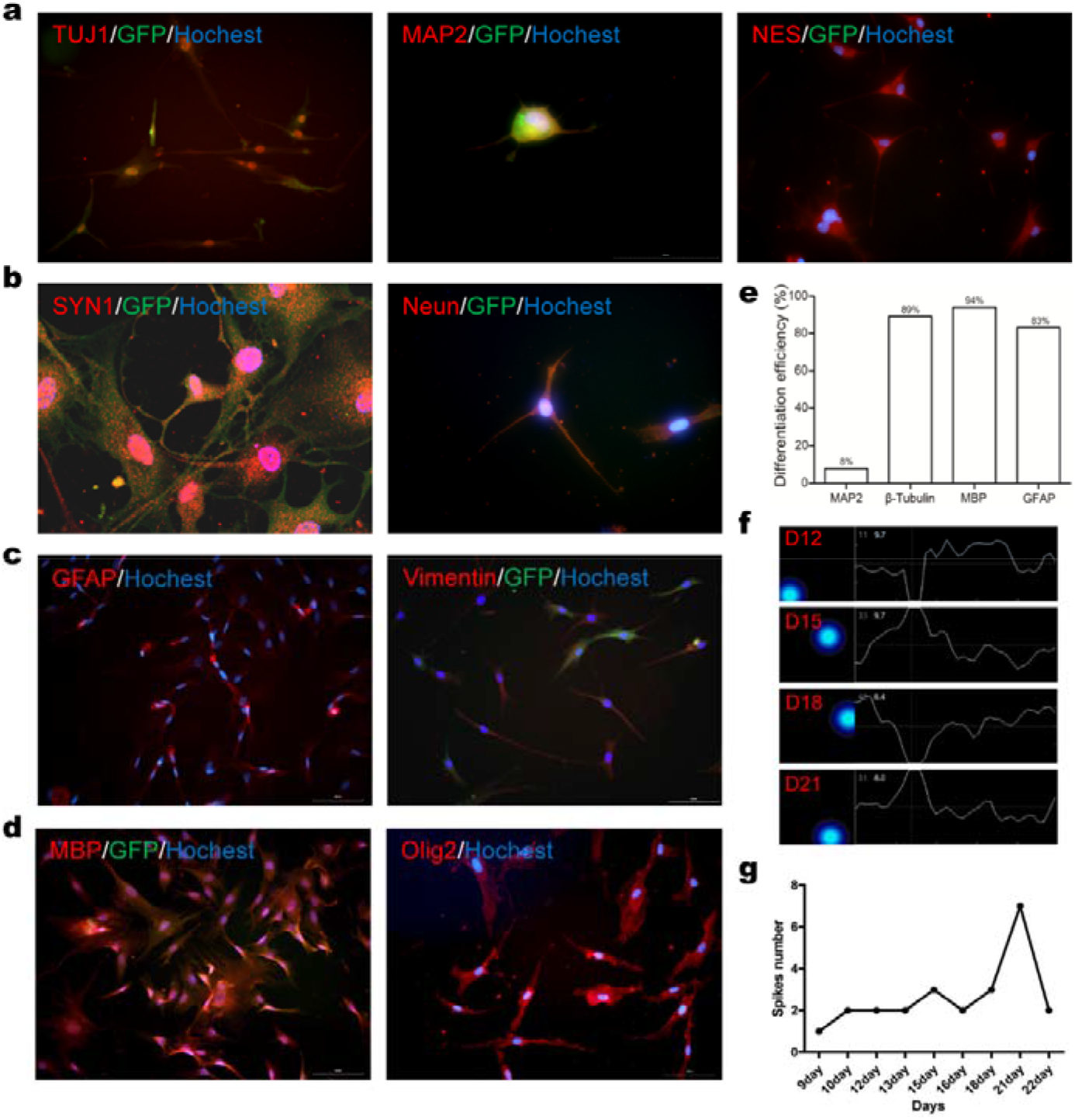
Multipotency of iNSCs In Vitro.a) hiNSC could be differentiated into Tuj1+, MAP2+ and NSE+ neurons by day 8 in culture after growth factor withdrawal. b) hiNSCs were able to differentiate into SYN1+ and NeuN+ neurons.c) hiNSC differentiate into subtypes of astrocytes, including GFAP+ and Vimentin+ positive cells. d) hiNSCs can robustly generate MBP+ and Olig2+ oligodendrocytes by 22 days in vitro. Scale bars represent 100 μm in (a)-(d). e) Quantification of Map2 +, β-Tubulin +, MBP+ and GFAP+ positive cells rate differentiated from hiNSCs.f) Cumulative activity map and spiking amplitude changes of differentiated neuron derived from hiNSCs at D12, D15, D18 and D21 using the MEA detection.

To study the electrophysiological function of neurons derived from iNSCs, we use microelectrode array (MEA) analysis to detect electrical activity in cell populations for 8–21 days, a technique based on the level of the electrical activity of the population. From day 9, the machine recorded spontaneous action potential. The number of spontaneous action potentials increased gradually and reached a peak on day 21, and then decreased on day 22. This trend continues until cell death (Figure 2f). The neurons differentiated from iNSCs show an obvious spontaneous firing pattern of the synchronous cluster which indicates that the neural network has matured. After excitation, the differentiated cells show even more and stronger excitation action potentials (Figure 2g, Figure S9, Supporting Information). Besides, during neuronal differentiation, synapsin-positive expressing cells can be observed, indicating that they can form synaptic connections between neurons ( Figure 2b), indicating the formation of synaptic connections between differentiated mature neurons. Thus, mir-302a-reprogrammed hiNSCs can differentiate into mature and functional neurons.

To assess the developmental and differentiation capacity of iNSCs in vivo, the hiNSCs, and miNSCs were microinjected into the cortex and hippocampus of 3 to 5 weeks of nude mice. Mouse and human-induced neural stem cells were detected by immunofluorescence near the transplantation site 2-4 weeks after transplantation. Transplanted hiNSCs and miNSCs can differentiate into NeuN and Tuj1 positive neurons, and GFAP, S100b positive astrocytes, MBP, and Olig2 positive glial cells (Figure 3a-f, Figure S10 and S11, Supporting Information). Therefore, mir-302a-iNSCs can not only survive but also differentiate into neurons, astrocytes, and oligodendrocytes in vivo. There was no teratoma formation within the mice brain after iNSCs transplantation. We also tested multiple iNSCs from different passages, with no teratoma formation in hippocampi. These observations indicated that mir-302a-hiNSCs not only possess the ability to survive, differentiate, and integrate into the adult brain but also not form teratoma in vivo.

**Figure 3.**
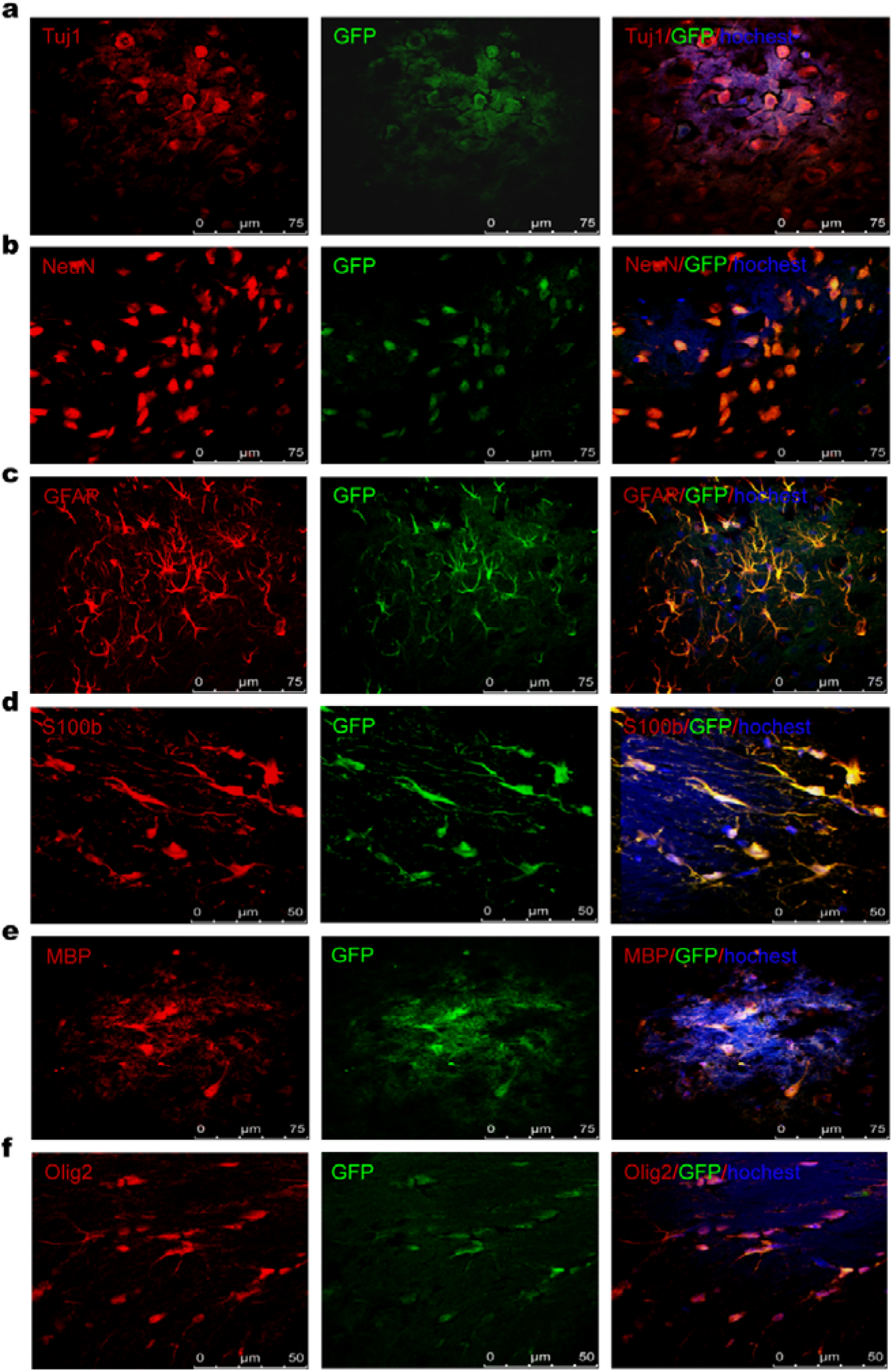
Mulitipotency of mir-302a-reprogrammed iNSCs in vivo. The mir-302a-reprogrammed iNSCs were injected into the striatum of 2-month-old nude mice. 3 weeks after post-transplantation, GFP-positive iNSCs migrated and integrated into the mice brain. a-b) Immunostains reveal that hiNSCs can differentiate into Tuj1+and NeuN+ neurons. c-d) GFP+ iNSC can differentiate into GFAP+and S100b+ astrocytes. e-f) Injected GFP+ cells co-expressing the oligodendrocytes cell marker MBP and Olig2. Scale bars represent 100 μm in (a) and 75μm in (b).

In summary, we provide an unprecedented method for rapidly and efficiently generating a Large number of high-purity iNSCs from human and mouse fibroblasts. At present, neural stem cells are reprogrammed by transcription factors or in combination with small molecules^[3, 25]^. MicroRNAs are often employed to enhance reprogramming efficiency by assisting transcription factors^[21]^. In contrast, our study demonstrates that a single mir-302a was sufficient to reprogram human and mouse fibroblasts efficiently to generate iNSCs.In contrast, our study demonstrates that a single mir-302a was sufficient to reprogram human and mouse fibroblasts efficiently to generate iNSCs. This result is completely different from the conclusion of previous studies that mir302 clusters must be used in combination and also with mir-367 to have reprogramming capabilities^[22]^. This provides a new proof of concept of the role of a single component of microRNA in the development and reprogramming of neural stem cells. Besides, because microRNA is only expressed in the cytoplasm and not in the genome, the use of mir-302a can avoid integration into the iNSC genome, eliminating the possibility of potential tumorigenicity and providing safety assurance for future cell therapy. The existing protocols that use translational factor or small-molecule compounds to reprogram somatic cells into iNSCs are tedious, requiring up to more than 2 weeks with relatively low reprogramming efficiency (0.08%-66%) and colonies formation efficiency ( 0.0002%-0.6% ). Instead, We have developed a method similar to that used by other researchers to grow neural stem cells on glass coverslips^[3]^, while reprogramming somatic cells with a single microRNA and small molecules to make neural stem cells, with a reprogramming efficiency of up to 90% within 3 days, the highest clone formation efficiency ( 3.04% ) of any date methods can be achieved. Moreover, the ability to produce large numbers of pure iNSCs from people of different ages and body parts provides the scalability necessary for developmental research, disease model making, drug screening, and large-scale cell therapy.

## Experimental Section

The experimental procedures and materials used are included in the Supporting Information

## Supporting information

supplemental methods,table and figuresa

## Supporting Information

Supporting Information is available from the Wiley Online Library or from the author

## Acknowledgements

Y.L , J.S and T.X contributed equally to this work. We thank Liqin Ren for animal feeding, Renhui Zhan for gene detection, Jing Guo and Tingting He for primary cell isolation from human eyelid and foreskin, Yanwei Wang for primary cell collection from human foreskin. The authors would like to thank Prof. Mengqing Xiang and Dr. Dongchang Xiao (Sun Yat-sen University) for providing the human and murine neural stem cells. Xuemei Hu for partial financial support and advice. This work was supported by the Shandong Province Natural Science Foundation Grants ZR2018LC008 (Y.W.), Yantai Science and Technology Innovation Development Plan 2020XDRH106 ( Y.W. )and the Fundamental Research Funds for the Central Universities 31020200QD020 (B.D.) of China.

## Conflict of Interest

The authors declare no conflict of interest.

## Data Availability Statement

The data that support the findings of this study are available from the corresponding author upon reasonable request

## Notes

### Competing Interest Statement

The authors have declared no competing interest.

