## supplemental methods,table and figuresa for "Efficient and Rapid generation of anti-aging neural stem cells by direct conversion Fibroblasts with A Single microRNA"

### Experimental Methods

**Human Samples.** Fresh human foreskin samples from two healthy male donors (at age of 26 and 29) were collected from Yuhuangding Hospital at Yantai, and fresh human eyelid samples from a healthy female donor (at age of 41) were collected from Chang'an Hospital at Xi'an. All the procedures were performed according to an Ethics Committee-approved protocol of Binzhou Medical University. The primary foreskin or eyelid cells were cut into pieces with ophthalmic scissors,enzymatically disassociated with (trypsin,0.25%,30-60min),and cultured in fibroblasts culture medium, DMEM-High Glucose with 1% penicillin/streptomycin,2 nM L-glutamine, and 10% FBS.

**Human iNSCs generation.** Human foreskin or eyelid derived fibroblasts were cultured on gelatin-coated glass coverslips at a density of $1x10^4$ cells/well in a 24-well-plate in fibroblast culture medium overnight. On the second day, the medium was changed into fresh medium containing mir-302a virus with polybrene and VPA. 24 hours post-transfection, Vitamin C were added within NSC induction media 500 μL (DMEM/F12 + 2% B27 +20 ng bFGF +10 ng/mL EGF + 1% DOX +1% L-glutamine +1% penicillin/streptomycin). After 72h, Puromycin was added to remove untransfected cells, then the media were changed into NSC culture media (DMEM/F12 +2% B27 +20 ng/mL bFGF+20 ng/mL EGF+1% L-glutamine+1% penicillin/streptomycin.). The medium was changed every 2 days. From 24h after transfection, some fibroblasts morphology changed, and a few small colonies formed and expand.

**Real-time PCR.** Total RNA was isolated from iNSCs with the TaKaRa kit (CatalogRR820A), RT-PCR was performed with The Light cycle 96 Real-Time PCR System (Roche). The PCR-program: 95 °C 30 s, 95°C 5s, 60°C 30s. Steps 2-3 were repeated 40-45 times. GAPDH gene was used as an internal control. Primers are listed as followed (F, Forward; R, Reverse; sequences are from 5' to 3'):

Nestin-F: CTGCTACCCTTGAGACACCTG;

Nestin-R: GGGCTCTGATCTCTGCATCTAC

Sox2-F: GTGAGCGCCCTGCAGTACAA;

Sox2-R: GCGAGTAGGACATGCTGTAGGTG;

Pax6-F: GTCTTCAAGCAACAACAGCAGCA;

Pax6-R: CGGATGCTGTCCACTCTCACAATA;

Oct4-F: GCTGGATGTCAGGGCTCTTTG;

Oct4-R: TTCAAGAGATTTATCGAGCACCTTC

**Differentiation of hiNSC.** Astrocytes differentiation: First, a 12-well-plate was coated with 40 μg/mL poly-lysine for 1 hour at 37ºC, and washed with PBS for 3 times. Then hiNSCs were plated onto those polylysine-coated wells, astrocytes culture medium (DMEM/F12 medium supplemented with 10%FBS, 1x NEAA, and 2 mM L-glutamine) was applied after hiNSCs attached to the bottom. Oligodendrocytes differentiation: hiNSCs were enzymatically disrupted into single cells and seeded onto a polylysine-coated glass coverslip and cultured in DMEM/F12 with 1×N2, 10 ng/mL PDGF and 3μM forskolin, half of the medium was changed every 2 days. 4 days later, forskolin was substituted with 200 mM Vc and cultured for 7 days. Neuron differentiation: For the generation of terminally differentiated neurons from hiNSCs, the cells were seeded at a density of 1 x $10^4$ cells per 12 mm glass coverslip coated with poly-ornithine/laminin in the neural induction medium (Neurobasal medium supplemented with 2% B27、500μM dbcAMP、10μM SB431542、10 ng/mL BDNF、10ng/mL NT-3、1μM Ken 、2 mM Glutamax-ITM and 1% penicillin/streptomycin. ), half of the medium was changed every 2 days.

**Immunofluorescence staining.** Cells were washed 3 times with PBS and then fixed with 4% paraformaldehyde (PFA, Sigma-Aldrich) for 10 min. Then cells were punched with 0.3% Triton for 10 min and blocked by 3% goat serum for 30 min at37°C. After that cells were incubated with primary antibodies overnight at 4 °C followed by secondary antibody staining for 45 min at 37°C. After nuclear staining with Hoechst 33342(Sigma-Aldrich), cells were observed and captured by Olympus CK51 microscope.

**In vivo transplantation assay.** 3 weeks wild-type C57BL/6 mice were anesthetized and fixed on a stereotaxic frame.2 μL P3 hiNSCs cell suspension was microinjected into one side of the telencephalon (x：-0.22 cm y：-0.26 cm z：-0.14 cm) for 10 min using a micro-syringe. After the injection, the needle is pulled after 5 min, and we pulled the needle out for 3 minutes. 2 to 3 weeks after transplantation, immunofluorescence detected the frozen section of the brain. All the procedures were performed according to an Ethics Committee-approved protocol of Binzhou Medical University.

**MEA recording.** For neuronal Spontaneous action potential analysis on MEA plates, 24-well MEA plates containing 16 electrodes each were coated with 50 μg/mL laminin for 3 h at 37 °C. The neurons derived from the differentiation of iNSCs were digested and centrifuged, the supernatant was removed, and the proper culture medium was added and mixed. 30 μL of the cell suspension to each pore was added to ensure that all drops apply to the electrodes for 1 h at 37 °C（25,000 cells in a well），then add 250 μL of culture medium to each well and then 259 μL of culture medium, changing the media by half every three days. DIV 9 to 21-day-old MEA cultures were used for analysis. 5 min recordings/1 h were performed every day except the day of media change using an MEA system (Axion Biosystems). We set the action potential spike detecting threshold to 5.5

standard deviations.

**Statistical information.** In general, data for each result were obtained from three sample. results are presented as the mean±SD.Statistical analysis was performed using the Microsoft Excel computer programs and sigma 6.0. Unpaired two-tailed Student’s t-tests and/or Mann–Whitney U-test were used to performe the difference in real time PCR results. $P < 0.05$ was considered statistically significant.

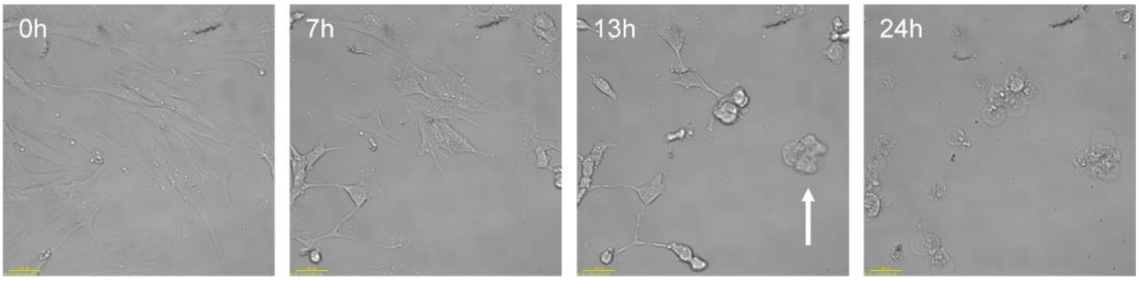


**Figure S1.** Formation of the first neurosphere from reprogrammed human fibroblasts transfected with mir-302 lentivirus.13 hours after transfection of human skin fibroblasts with mir-302a, the first early neurosphere-like colonies generated in bright-field image with Living cell workstation. After 24 hours, the vast majority of cells become neurospheres-like colonies. （Deltavision Elite, GE ）. Scale bar: 40 μm, T represents time.

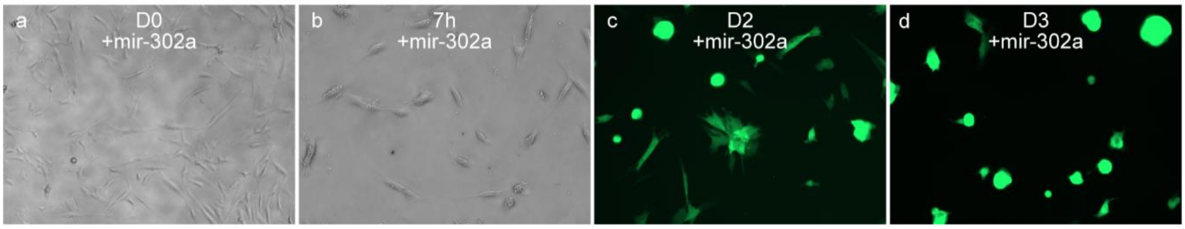


**Figure S2.** Generation of hiNSCs from human eyelid fibroblasts (HEFs). **(**a-b), Morphological changes of HEFs infected with mir-302a within 7 hours. c-d, Numerous cell GFP -positive clusters could be seen at 2 and 3 days after transfection. From day 2, many GFP-positive neurosphere-like structures formed. On day 3, it reprogrammed most cells to form GFP-positive neurosphere-like structures.

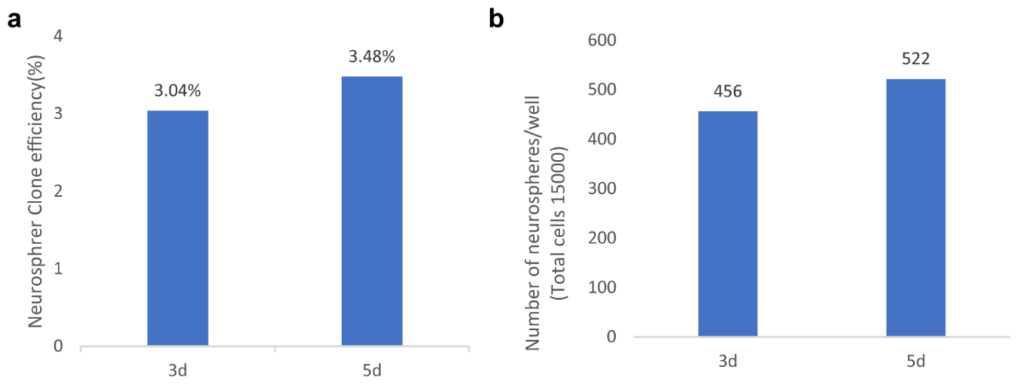


**Figure S3.** Cloning efficiency of hiNSCs from human Eyelid fibroblasts. Quantification of neurospheres induced by mir-302a with Eyelid fibroblasts. HEFs (1.5 × 104) were seeded into each well of 24-well plates, infected with mir-302a lentiviruses Neurospheres in each well were counted at day 3 and 5 following virus infection. a) Cloning formation efficiency of HEFs cells from 41-year-old female donors described above at day 3 and 5. b) Quantification of neurospheres induced by mir-302a with HEFs on day 3 and 5 after transfection.

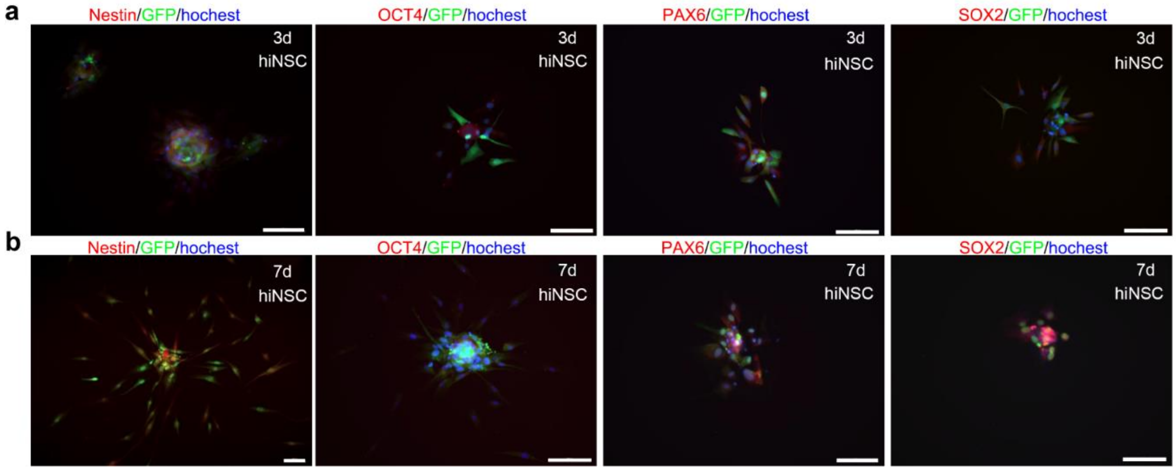


**Figure S4.** Characterization of hiNSCs from Eyelid fibroblasts. Neurospheres induced from HEFs by mir-302a on day 3 and 7 were immunoreactive for Nestin, Oct4,Pax6 and Sox2. a) On day 3, most reprogrammed cells then expressed the neural stem cell markers Nestin, Pax6, and Sox2. b) Nestin, Pax6, and Sox2 markers had increased on day 7. It expressed Oct4 on day 3 and decreased on day 7.

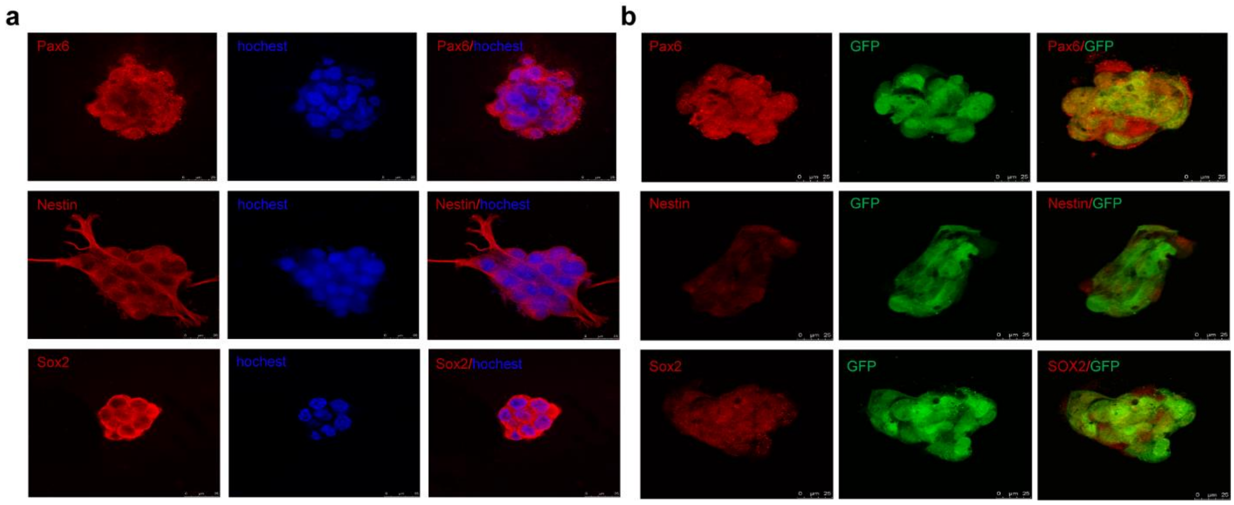


**Figure S5.** Immunostaining of the control NSCs (Ptf1a-induced hiNSCs) and mir-302a induced hiNSCs.a)The control HFFs infected with Ptf1a lentiviruses derived neurospheres were Pax6, Nestin, and Sox2 positive expression in Immunostaining. b) Neurospheres induced from HFFs by mir-302a were immunoreactive for Pax6, Nestin, and Sox2.

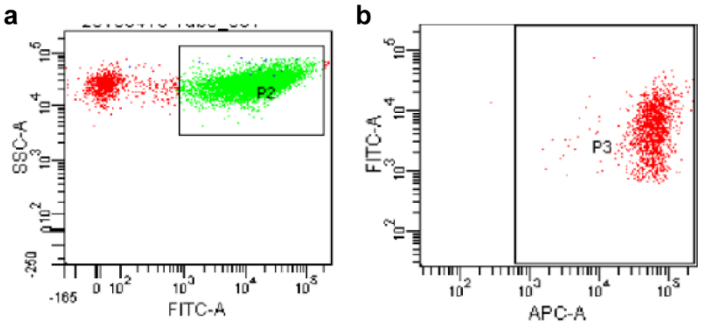


**Figure S6.** Flow cytometry measurements of the purity and yield of mir-302a-hiNSCs.a) Flow cytometry measurements of the yield of hiNSCs conversion showed that nearly 90% of surviving infected fibroblasts are Nestin positive on day 3. b) The purity of Nestin positive cells from these colonies were about 97%.

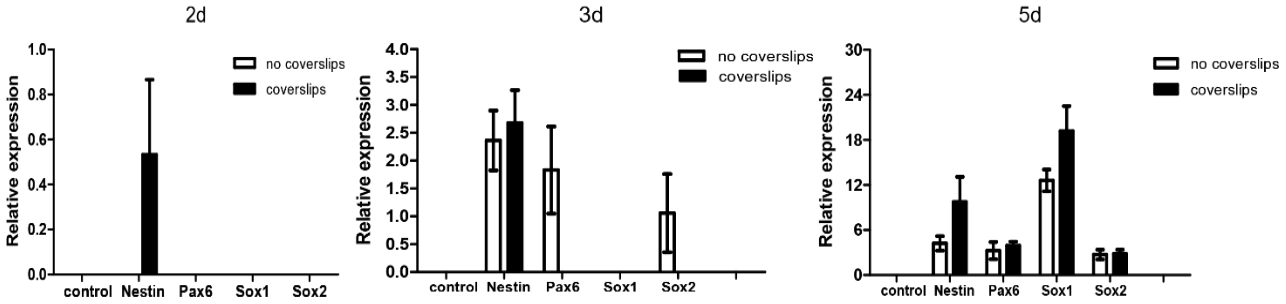


**Figure S7.** qRT-PCR analysis of mir-302a-hiNSCs.There was a significant difference in gene expression between direct cell seeding on plastic and on gelatin-coated glass coverslips methods to reprogram the HFFs into hiNSCs at different time point. There was a great increase in expression of Nestin, Pax6, Sox1, and Sox1 in hiNSCs generated on gelatin-coated glass coverslips at the same time, as well as on gelatin-coated plastic. As time increased, expression increased in both groups, and gene expression remained stronger on glass coverslips than on plastic plates.

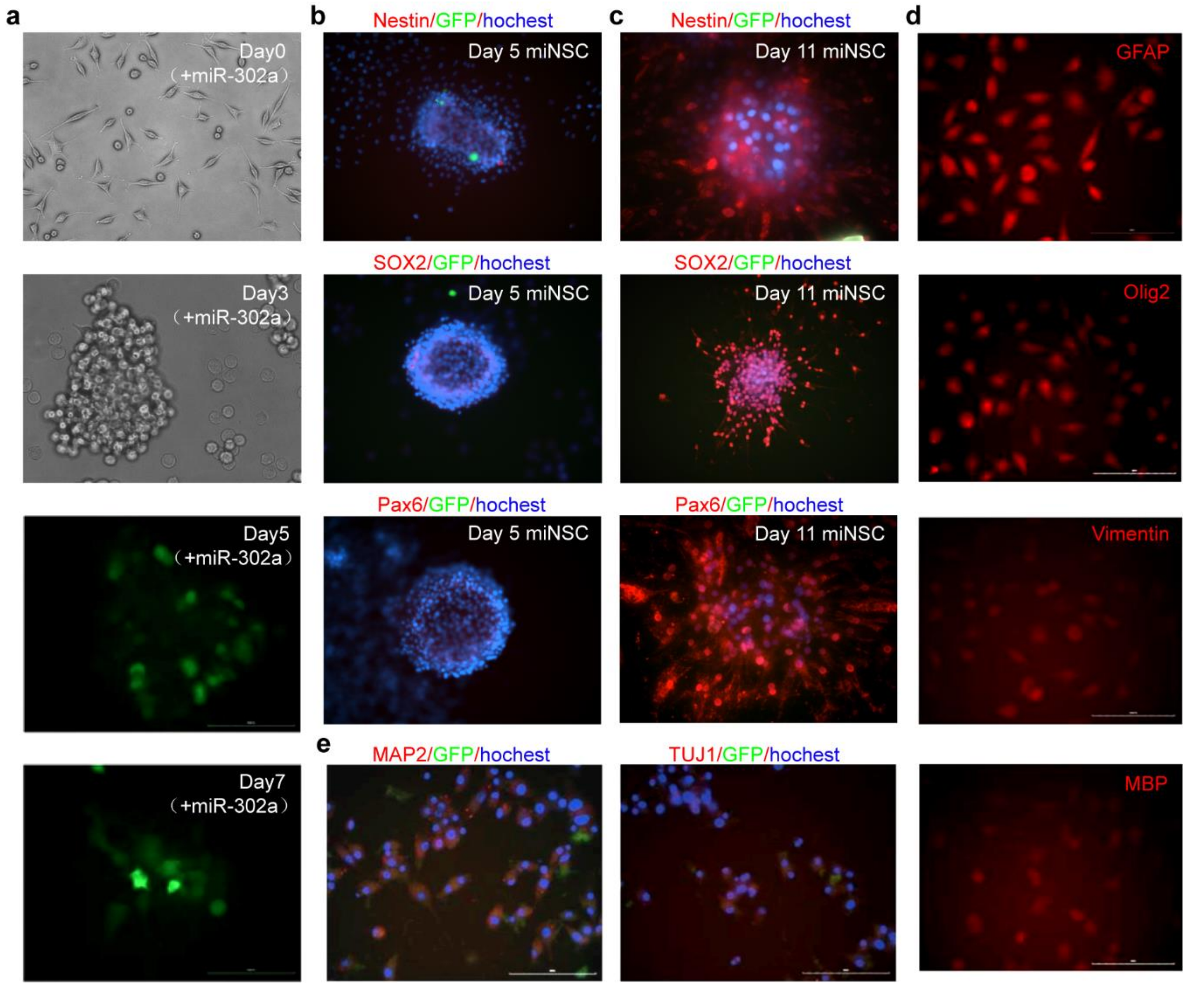


**Figure S8.** mir-302a reprograms mouse L929 fibroblasts into miNSCs.a) Representative fields show a time course of colony formation at days 0，3，5，and 7 after mouse L929 single-cell plating with mir-302a transfection.b) Immunofluorescent staining of mouse L929 fibroblasts-miNSCs for Nestin，Sox2，and Pax6 on day 5.c) Immunofluorescence staining for Nestin, Sox2, and Pax6 of miNSCs induced by mouse L929 fibroblasts produced at 11 days after secondary spheroid formation in culture. d) Representative images of in vitro differentiation of miNSCs into GFAP+, Olig2+, Vimentin+ and MBP+ astrocytes and oligodendrocytes.e)Representative images of in vitro differentiation of miNSCs into MAP2+ and Tuj1+ neurons.

Data shown are representative images from three independent experiments that gave similar results.

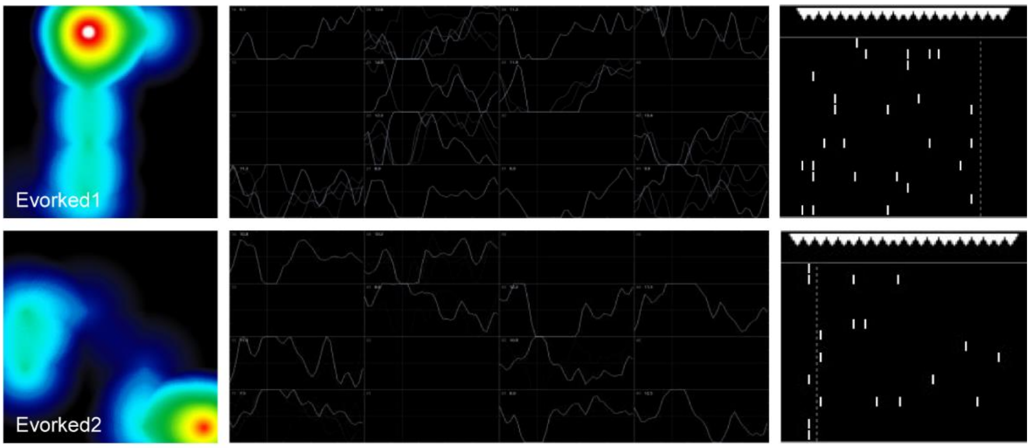


**Figure S9.** Evoked action potentials of differentiated neurons from mir-302a-hiNSCs.A large number of action potentials called evoked action potentials can be generated by electrical stimulation and displayed by action potential thermograms, amplitudes, and raster images. The firing synchronism, firing frequency, and the number of active electrodes of the differentiated neurons were increased.

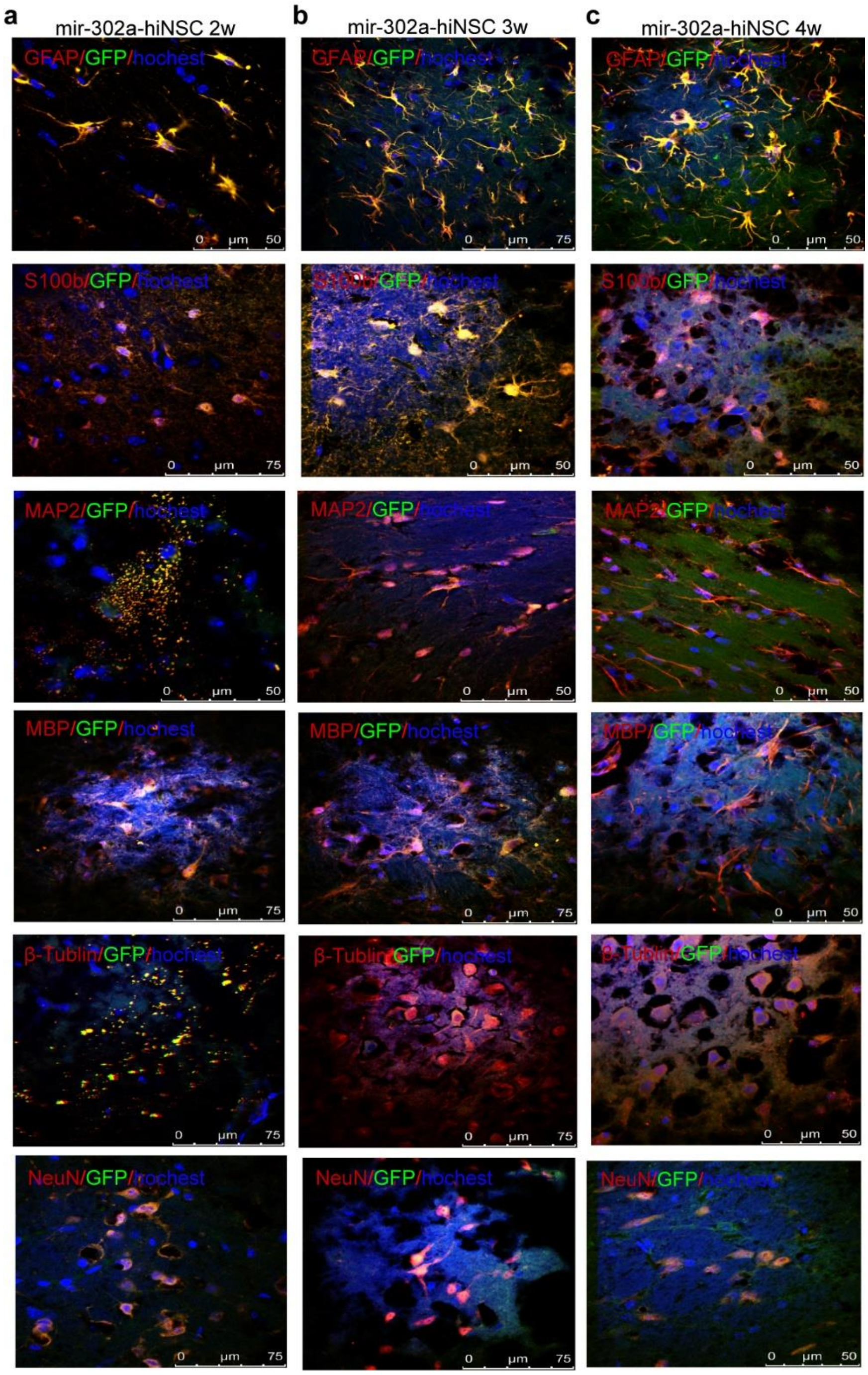


**Figure S10.** Mulitipotency of mir-302a-reprogrammed hiNSCs in vivo. The mir-302a-reprogrammed hiNSCs were injected into the striatum of 2-month-old nude mice. 2,3 and 4 weeks after post-transplantation, GFP-positive iNSCs migrated and integrated into the mice brain.a) Immunostains reveal that hiNSCs can differentiate into GFAP+ and S100b+ astrocyte, Tuj1+and NeuN+ neurons, and MBP+and olig2+ oligodendrocytes at 2 weeks after post-transplantation.b) Immunostains reveal that hiNSCs can differentiate into astrocytes, neurons, and oligodendrocytes at 3 weeks after post-transplantation.c) At 4 weeks after post-transplantation. GFAP+ and S100b+ astrocyte, Tuj1+ and NeuN+ neurons, and MBP+ and olig2+ oligodendrocytes derived from hiNSCs could be detected by Immunostains.

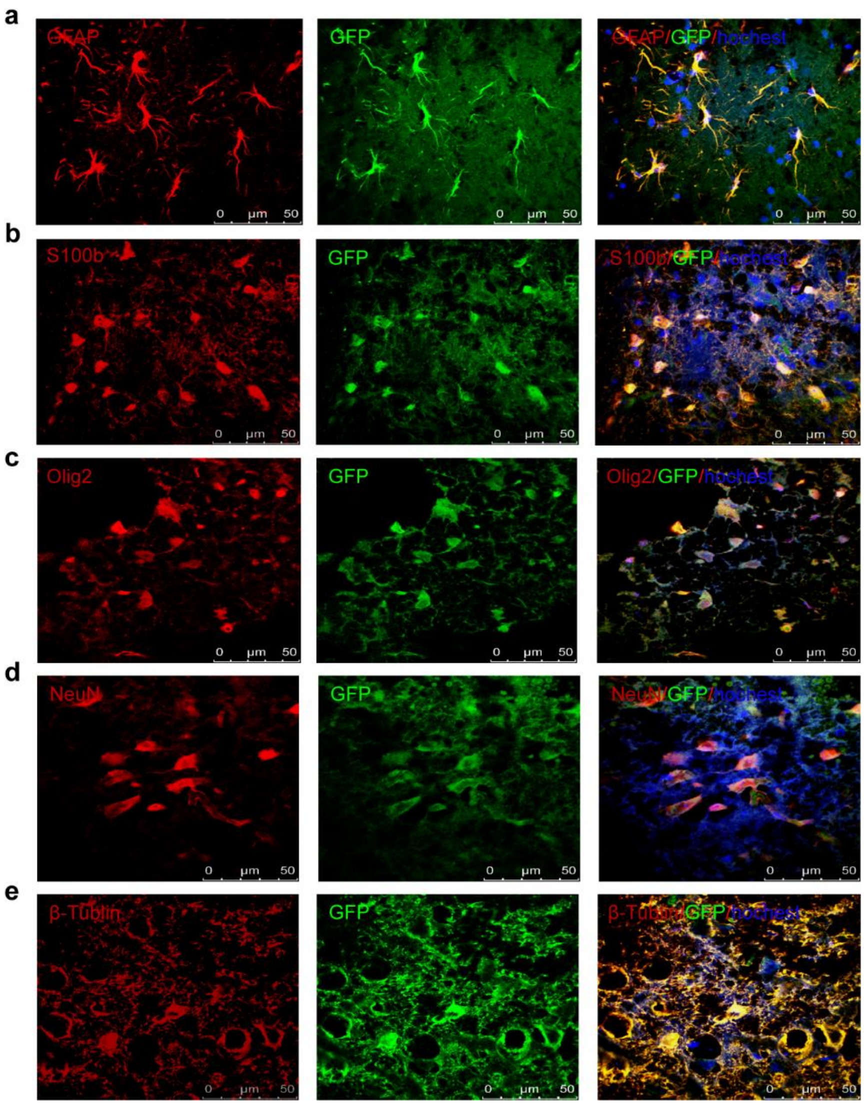


**Figure S11.** Differentiation potential of mir-302a-reprogrammed miNSCs in vivo. The mir-302a-reprogrammed miNSCs were injected into the striatum of 2-month-old nude mice. 3 weeks after post-transplantation, GFP-positive miNSCs migrated and integrated into the mice brain. a-b) Immunostains reveal that miNSCs can differentiate into GFAP+and S100b+ astrocytes. c) GFP+ miNSC can differentiate into Olig2+ oligodendrocytes. d-e) Injected GFP+ cells have the neuron cell marker NeuN and Tuj1.

**Table S1.** Overview of direct conversion methods for the derivation of human iNSCs.

| Cell source chrom | Transcription Factor（s） | Others supplements / Pharmacological compound | Conversion efficiency | The earliest time of neurospheres emerge | Mean time to neurospheres formation | Clonal formation efficiency | Time of appearance of electrophysiological action potential | Reference |
|---|---|---|---|---|---|---|---|---|
| Human fibroblasts | **mir-302a** | | **86.8%** | **13 hours** | **2-3 day（d）** | **3.04% （3 d）；8.02%（7 d）** | **10 day（d）** | **This submission** |
| Mouse fibroblasts Human fibroblasts | SOX2 | | | 4 d | 6-10 d | | | Ring KL et al. cell stem cell,2012 |
| Human fibroblasts | SOX2 | | | 48 hours | 4 d | | | Bagó JR et al. Sci Transl Med, 2017 |
| human urine cells | OCT4, SOX2, SV40LT, KLF4 | mir-302/367 | | 12 d | 25 d | 0.2% | | Wang L et al. nature methods, 2013 |
| hDFs | SOX2 | let-7b，HMGA2 | | | 7-20 d | 0.2%-0.6% | | Kyung RY et al. Cell Reports，2015 |
| Mouse fibroblast and human foreskin fibroblasts | Ptf1a | | | | | | 2 weeks | Xiao D et al. Nat. Commun,2018 |
| Cord Blood CD34+cells | OCT4 | | | | | | | Yang H et al. stem cells translational medicine,2015 |
| Adult human fibroblasts | OCT4 | | | 7+14 d | 3-4 weeks | | | Ryan R. M et al. Stem Cells,2014 |
| Human Neonatal and Adult Blood Cells | OCT4 | SMAD+GSK-3、(SB431542, LDN-193189, Noggin, CHIR99021) | | 8–10 d | | 0.024% | | Lee JH et al. Cell Rep, 2015 |
| human neonatal (foreskin) fibroblasts (HNFs | Zfp521 | BAM groups | | 24 d | 24-42 d | 0.4-0.7% | | Shahbazi E et al. Stem Cell Reports,2016 |
| human UCB-MSC | SOX2 | | 25% | 14 d | | 0.015% | | Kim B et al. Cell Transplant,2018 |
| Human fibroblasts from AD patients，healthy person | SOX2 | 9 small molecules | 0.038%-0.091 % | 12 d+6-8 d | 18-20 d | 0.09% | | Liu Y et al. J Alzheimers Dis, 2020 |
| Human Fibroblasts | OCT4 | RM | 0.94% | | | | | Ryan M et al. stem cells,2014 |
| adult human peripheral blood cells (PBCs) | SOX2，c-myc | CHIR99021、A83-01、hLIF、Trany1 | | 10 d | 10-21 d | 0.08% to 0.66% | 8–12 weeks | Sheng C et al.Nat. Commun,2018 |
| Human Dermal Fibroblast | c-MYC，SOX2 | | 0.2%–0.5% | | | | | Daekee K et al. Mol Ther Nucleic Acids, 2019 |
| human neonatal dermis-derived fibroblasts or adult adipose-derived stem cells | OCT4, KLF4, SOX2, c-MYC | | 0.07%(neonatal HFF) 0.01%(adult HASC) | | 30-60 d | | 8 weeks | Dana M et al. Stem Cell Reports,2016 |

| | | | | | | | | |
|---|---|---|---|---|---|---|---|---|
| human skin fibroblasts | OCT4, SOX2, Klf4, c-Myc | | | 17 d | | 0.20% | | Sandra M et al. Journal of Visualized Experiments,2015 |
| human fibroblast of amilial and Sporadic Parkinson’s Disease Patients | OSKM | | | 7 d | 18~21 d | | | Lee M et al. Int J Stem Cells,2019 |
| Postnatal and adult human and monkey fibroblasts | OSKM | LIF, SB431542, CHIR99021 | 0.03%–0.08% | 13 d | 20 d | | 10 weeks | Lu J, et al. Cell Rep,2013 |
| Human PBMNCs | Klf4, OCT3/4, SOX2,c-MyC | | | 12 d | | | | Zheng W et al. J Vis Exp,2019 |
| Human fetal fibroblasts | SOX2，c-Myc | Brn2 or Brn4 | | 5-7 d | 6-7 d | 20-60 colonies / 10000 cells/well | 4–6 weeks | Qingjian Z, et al. The journal of biology chemistry,2014 |
| Human bone marrow cells | Msi1，Ngn2，MBD2 | | | | | | | Vonderwalde I et al. Transl Stroke Res. 2020 |
| Human fibroblasts | SOX2, HMGA2, BRN4, SKM+SV40LT (BSKMLT) | phorbol-12-myristate-13-acetate, CHIR99021, SB431542 | | 7 d | 2 weeks | 1700 colonies/4 weeks | | Kwak TH et al. Int J Stem Cells,2020 |
| PBMC、ADFs、FPFs | BRN2, SOX2, KLF4, MYC, TLX, ZIC3 | BKSZ+CAPT | | 14 d | 19-24 d | 0.015-0.166% (19 d) | 10 weeks | Thier MC et al. Cell Stem Cell,2019 |
| human urine-derived cells | OCT4, SOX2, KLF4, GLIS1 | Purmorphamine, Forskolin, Vitamin C, Sodium butyrate (N) | | 8 d | 8-12 d | 11clonies/100,000cells | 21 d | Kang PJ et al. Cells, 2019 |
| Human Fibroblasts | 25nTFs,15TFs,13Tfs,6TFs,7TFs | | 10.54-11.22% | 6 d | | | 4-5 weeks | Pei SH et al. Stem Cell Reports,2017 |
| human fibroblasts | OCT4, or OCT4/SOX2/KLF4/p53shRNA | SB431542 and CHIR99021 | | | 2-3 weeks | | | Saiyong Z et al. nature protocols,2015 |
| Human bone marrow-derived cells, foreskin fibroblasts, keratinocytes | Msi1, Ngn2, and MBD2 | | 72 ± 8.97% and 86.25 ± 7.23% | 12 d | 2 weeks | | | Ahlfors JE et al. Stem Cell Res Ther. 2019 |
| adult human peripheral blood mononuclear cells | OCT4, SOX2, NANOG, LIN28, c-YC, KLF-4 and SV40LT | | | 10 d | 30 d | | 7 weeks | Xihe T et al. Stem Cell Research,2016 |

**Table S2.** Overview of direct conversion methods for the derivation of mouse iNSCs.

| **Cell source** | **Transcription Factor （s）** | **Others supplements / Pharmacological compound** | **Conversion efficiency** | **The earliest time of neurospheres emerge** | **Mean time to neurospheres formation** | **Clonal formation efficiency** | **Time of appearance of electrophysiological action potential** | **Reference** |
|---|---|---|---|---|---|---|---|---|
| Rat fibroblasts and astrocytes | Zfp521 or sox2 | | 46.1 ± 2.9% | | | | 6 weeks in vivo | Zarei-Kheirabadi M,et al .Stem Cell Res Ther,2019 |
| Mouse fibroblasts Human fibroblasts | SOX2 | | | 8 d | 6-10 d | 0.13%-0.96% | 21 d | Ring KL et al. cell stem cell,2012 |
| Mouse fibroblast | Ptf1a | | | 6 d | 9-14 d | 0.5% at 14 day | 3 weeks | Xiao D et al. Nat. Commun,2018 |
| Mouse astrocytes | Zfp521 | | 41.7 ± 6.2% | | | | 60 d | Zarei-Kheirabadi M, et al. J Cell Physiol. 2019 |
| MEFs | Sox2, Klf4, c-Myc, Oct4 | | | 11 d | 19 d | 11 neurosphere /130,000 cells | 3 weeks | Thier M, et al. Cell Stem Cell,2012 |
| MEF | Oct4, Sox2, Klf4, c-Myc | | 0.07% | 11 d | 11-15 d | 0.69%-0.5% | 20 d | Kim J, et al. PNAS,2011 |
| mouse fibroblasts | Sox2, Klf4, c-Myc, Brn4 | | | 4-5 week | | | | Kim SM, et al. Nature protocols,2014 |
| mouse fibroblasts | BSKM. Brn4, Sox2, Klf4, c-Myc (BSKM) | | 6.3 ± 0.43% | 4 weeks | 4-5 weeks | | 14-16 d | Kim SM, et al. J. Biol. Chem,2016 |
| mouse fibroblasts | Brn4,Sox2,Klf4,c-Myc, plus E47/Tcf3 | | | 4-5weeks | 4-5weeks | 1-5 clusters/5 × $10^4$ cells | 7-16 d | Han DW, et al. cell stem cell,2012 |
| C57BL/6MEFs; C3H MEFs | Brn4,Sox2, Klf4,c-Myc (BSKM) | | 2.99%(C57BL/6MEFs) 8.22%(C3H MEFs) | | 6 week | | | Kim SM, et al. Stem Cell Research,2016 |
| mouse fibroblasts | Ezh2,Jarid2,Mtf2,Nanog,Pou5f1,Sall4,Smarca4, Sox2, Suz12, Tcf3 | | | | | | | Yaqubi M, et al. Stem Cell Research & Therapy,201 |
| mouse fibroblasts,Liver Cells,B Lymphocytes | Brn2,Hes1,Hes3,Klf4, Myc,Notch1,(NICD),PLAGL1, Rfx4 13TFs；14Tfs | | 1.00% | | 30 d | | 4 weeks | Cassady JP, et al. Stem Cell Reports, 2014 |

| | | | | | | | | |
|---|---|---|---|---|---|---|---|---|
| mouse fibroblasts | Oct4,Sox2,Klf4,c-myc, Brn2,FoxG1,et al.11 factor | | 12.3 % | 24 d | 25 d（13d FoxG1+Sox2） | 3-317 colonies/20000 cells | 25 d | Lujan E, et al. PNAS, 2011 |
| Astrocytes mouse | SOX2 ASCL1 | | 23.2%± 5.3% | | | | | Niu W, et al. Stem Cell reports,2015 |
| sertoli cells | Sox2,Pax6, Ngn2, Hes1， Id1, Ascl1, Brn2.c-Myc, and Klf4 | | 0.70% | | | 0.0002% | 3 weeks | Chao S, et al. Cell Research,2012 |
